# Pasakbumin A controls the growth of *Mycobacterium tuberculosis* by enhancing the autophagy and production of antibacterial mediators in mouse macrophages

**DOI:** 10.1101/348144

**Authors:** Hyo-Ji Lee, Hyun-Jeong Ko, Seung Hyun Kim, Yu-Jin Jung

**Affiliations:** Department of Biological Sciences and Institute of Life Sciences, Kangwon National University, Chuncheon, 24341, Republic of Korea; College of Pharmacy, Kangwon National University, Chuncheon, 24341, Republic of Korea; College of Pharmacy, Yonsei University, Incheon, 21983, Republic of Korea

**Keywords:** Pasakbumin A, Antimycobacterial activity, Autophagy, Host-directed therapy (HDT), *Mycobacterium tuberculosis*

## Abstract

Tuberculosis (TB) is a chronic infectious disease caused by *Mycobacterium tuberculosis* (Mtb) and remains a major health problem worldwide. Thus, there is an urgent need to identify new and more effective drugs to treat emerging multidrug-resistant TB (MDR-TB) and to reduce the side effects of anti-TB drugs, such as liver toxicity and other detrimental changes. In this study, to develop a novel candidate drug for effective TB treatment with few side effects in the host, we selected pasakbumin A isolated from *Eurycoma longifolia* (*E. longifolia*) Jack, which protected host cells against Mtb infection-induced death. Pasakbumin A significantly inhibited intracellular Mtb growth by inducing autophagy via the ERK1/2-mediated signaling pathway in Mtb-infected macrophages. We further investigated whether pasakbumin A could be used as a potential adjuvant for TB treatment. Treatment with pasakbumin A and the anti-TB drug rifampicin (RMP) potently suppressed intracellular Mtb killing by promoting autophagy as well as TNF-α production via the ERK1/2- and NF-κB-mediated signaling pathways in Mtb-infected cells. Our results suggest that pasakbumin A could be developed as a novel anti-TB drug or host-directed therapeutic (HDT) strategy to protect against host cell death and improve host defense mechanisms against Mtb infection in macrophages.

## Introduction

Tuberculosis (TB) is still one of the oldest known human diseases and a major cause of mortality among the infectious diseases [1]. *Mycobacterium tuberculosis* (Mtb), the causative agent of TB, is a highly successful facultative intracellular pathogen that can persist within host phagocytes[2]. Mtb infection usually begins after inhalation of aerosol droplets that contain bacteria into the pulmonary alveoli. After inhalation, Mtb is recognized by resident alveolar macrophages, dendritic cells and recruited monocytes through various pattern recognition receptors (PRRs)[3]. These receptors initiate diverse signal transduction pathways, including the nuclear factor-kappa B (NF-κB) and mitogen-activated protein kinase (MAPK) signaling pathways, which induce the production of cytokines and chemokines in host cells[4]. Induction of these effector molecules regulates bacterial growth and promotes the adaptive immune response. Mtb is also ingested by phagocytosis to form phagosomes containing Mtb-antigen (Mtb-Ag). After phagocytosis, mycobacterial antigens are processed and presented to Mtb-specific CD4^+^ T cells and CD8+ T cells, which produce several cytokines to activate macrophages and lymphocytes[5]. However, Mtb can survive and persist inside macrophages in the dormant stage for a long period by interfering with the host immune system to avoid elimination by the effector immune cells[6, 7].

Autophagy is a conserved lysosomal self-digestion process that involves turnover of cellular constituents to maintain cellular homeostasis[8]. This process also functions as an innate immune defense mechanism against infectious pathogens by fusing the lysosome with a double-membrane-bound autophagosome, which can sequester cytoplasmic material and pathogens[9, 10]. The autophagic process is tightly regulated by the action of autophagy-related (Atg) proteins, such as beclin-1 and microtubule-associated protein 1A/1B-light chain 3 (LC3)[11, 12]. Because cytosolic LC3 (LC3-I) is conjugated with phosphatidylethanolamine (PE) to form membrane-bound lapidated LC3 (LC3-II) during autophagy[13], the conversion of LC3-I to LC3-II is commonly used to measure and monitor autophagic events. However, Mtb secretes an enhanced intracellular survival (Eis) protein which inhibits autophagy and increases IL-10 expression[14]. Although many studies have shown that the activation of autophagy not only enhances phagosome-lysosome fusion but also regulates Mtb growth in host cells[15], Mtb has evolved several mechanisms to modulate or exploit the autophagic process [16–18].

Current TB treatment is based on multidrug chemotherapy. According to the WHO guide lines, a multidrug regimen for TB includes administration of first-line drugs consisting of rifampicin (RMP), isoniazid (INH), pyrazinamide (PZA), and ethambutol (EMB) for 2 months followed by INH and RMP for 4 months[19]. However, prolonged regimens using the same few drugs have resulted in poor patient compliance, which leads to the emergence of strains with resistant to available anti-TB drugs and the generation of multidrug (MDR) and extensively drug-resistant (XDR) Mtb[20–22]. Due to the increased emergence of drug-resistant Mtb strains, there in an urgent need for the development of new anti-TB drugs. Recently, attention has focused on a new and emerging concept in TB treatment known as host-directed therapy (HDT), which focuses on key components of host anti-mycobacterial effector mechanisms and limiting inflammation and tissue damage[23–25]. Therefore, in this study, we identified a novel anti-TB drug from natural compounds that exhibited antibacterial activity by enhancing host anti-TB effector mechanisms in mouse macrophages. To screen the anti-Mtb activities of selected natural compounds, we measured the bacterial growth in Mtb-infected macrophages after treatment with each compound. The best candidates among the selected natural compounds was pasakbumin A isolated from *Eurycoma longifolia* (*E. longifolia*) Jack, which is commonly used as a traditional herbal medicine to treat fever, malaria, ulcers and TB[26]. We observed significant inhibition of intracellular Mtb growth via induction of autophagy and increased production of NO and TNF-α in pasakbumin A-treated cells during Mtb infection. In addition, the combined treatment of the anti-TB drug RMP and pasakbumin A strongly reduced intracellular Mtb growth by promoting autophagy and inflammatory cytokine production via the ERK1/2- and NF-κB-mediated pathways in Mtb-infected cells. Together, our results suggest that pasakbumin A can enhance the Mtb-killing activity of macrophages by inducing autophagy. Pasakbumin A could be developed as an HDT drug and/or therapeutic strategy that enhances innate immune functions and modulates inflammatory responses for TB.

## Results

### Pasakbumin A controls intracellular Mtb growth by increasing the production of TNF-α and NO in Mtb-infected macrophages

Recent studies have shown that various compounds extracted from leaves, stem and root of *E. longifolia* exhibit potent antibacterial activity against pathogenic Gram-positive and Gram-negative bacteria[27]. However, the specific compounds extracted from E. *longifolia* that control intracellular Mtb growth remain poorly understood, although *E*. *longifolia* is also used as a traditional medicine for patients with TB in Malaysia. To identify the mycobactericidal activity of the compounds extracted from *E*. *longifolia* in macrophages, we assessed intracellular bacterial survival in H37Rv-infected cells treated with each compound. We obtained a specific compound, pasakbumin A, with potential mycobactericidal activities against Mtb infection. Pasakbumin A significantly suppressed intracellular Mtb growth in macrophages, although it was not directly toxic to Mtb (**Fig. 1A, 1E**). In addition, TNF-α production substantially increased in pasakbumin A-treated Raw264.7 macrophages during Mtb infection, but IL-10 production significantly decreased (**Fig. 1B**). Notably, we found that pasakbumin A strongly enhanced NO production and NOS2 expression in Raw264.7 macrophages during Mtb infection compared with those in Mtb-infected cells (**Fig. 1C**). Recently, Tousif *et al.* showed that the anti-TB drug INH strongly induces apoptosis of activated CD4^+^ T cells and reduces Mtb antigen-specific immune responses[28], suggesting that anti-TB drugs induce host cell damage as well as mycobacterial killing. In contrast to most commonly used anti-TB drugs, pasakbumin A was sufficient to protect Mtb-infected Raw264.7 cells compared to cells infected with Mtb alone, and it did not exhibit any cytotoxic effects in
macrophages (**Fig. 1D**). These results suggest that pasakbumin A controls intracellular Mtb growth by enhancing NO and TNF-α production by macrophages and protects against host cell death during Mtb infection.

**Fig 1.**
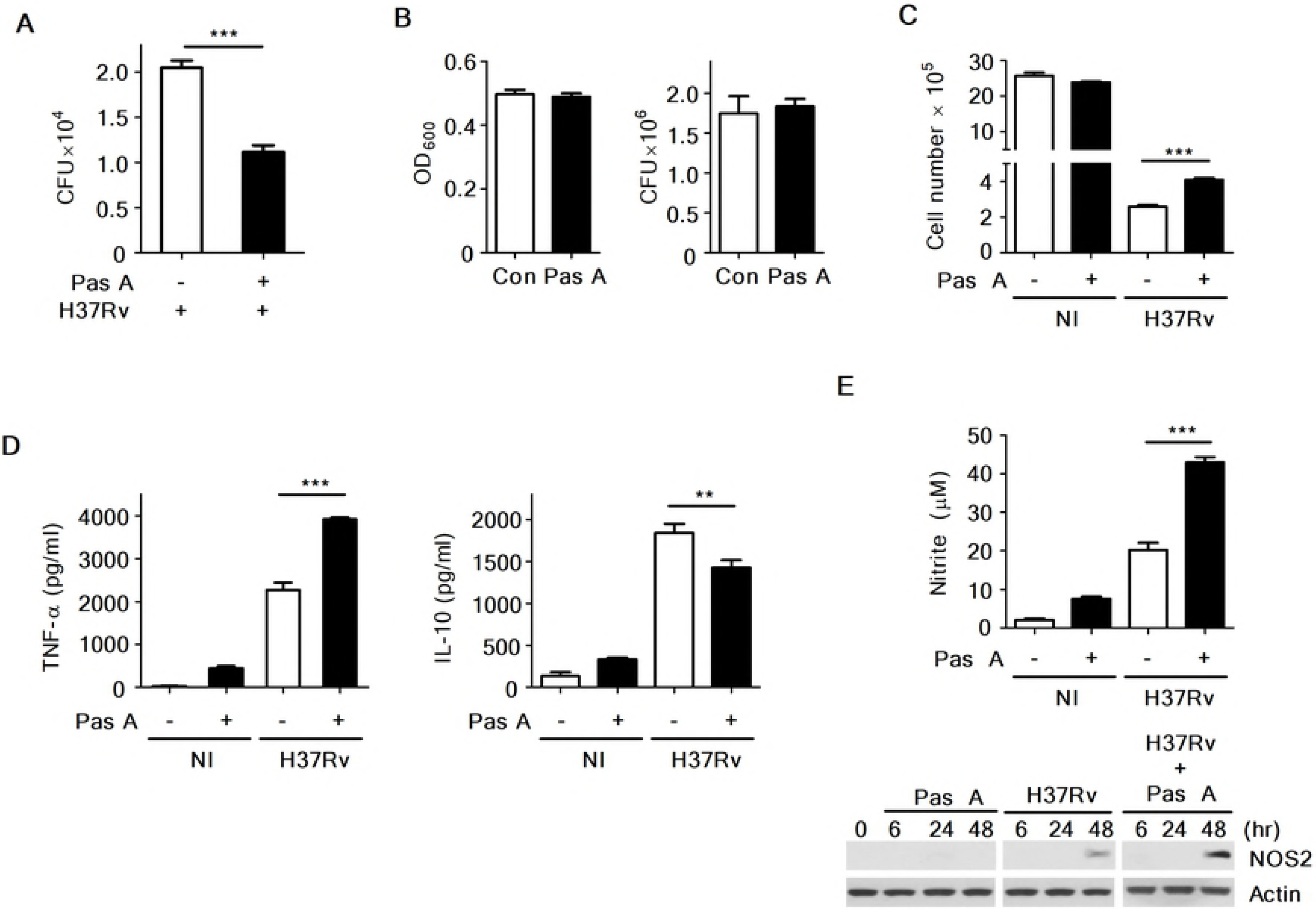
Pasakbumin A controls intracellular Mtb growth by increasing the production of NO and pro-inflammatory cytokine in H37Rv-infected macrophages. Raw264.7 macrophages were stimulated with pasakbumin A (Pas A, 10 μM) for 48 h after infection with H37Rv (at MOIs of 1 or 5). (A) Intracellular bacterial survival was determined by counting the number of CFUs at 3-weeks after inoculation. (B) TNF-α and IL-10 production in culture supernatants from non-infected (NI) or H37Rv-infected cells was measured by ELISA. (C) Bar graph in the left panel represents NO production (as indicated by the nitrite level) with the diazotization (Griess method) assay. Right panel represents NOS2 protein by western blot analysis. Actin served as a loading control. A complete western blot images is provided in **S1 Data File**. (D) Cell viability was assessed using a trypan-blue exclusion assay. (E) The cultures were grown in 7H9 medium supplemented with 10% ADC containing 0.2% glycerol at 37°C for 72 h with or without pasakbumin A (Pas A, 10 μM). Left, bacterial growth was measured as OD_600_ at 72 h after Mtb inoculation. Right, colony-forming units (CFUs) were measured by plating bacterial dilutions onto 7H10 agar supplemented with 10% OADC containing 0.5% glycerol at 72 h. Error bars represents the standard deviation of the mean. Statistical significance is indicated as **, *p*<0.01 and ***, *p*<0.001.

### Pasakbumin A activates the ERK1/2-mediated signaling pathway and induces autophagy in Mtb-infected macrophages

To examine the molecular mechanisms of pasakbumin A during Mtb infection, we investigated the intracellular pathway involved in the pasakbumin A-mediated response to Mtb via activation of various molecules associated with NF-κB and MAPK. The results indicated that the levels of phosphorylated ERK1/2, iκB-α and NF-κB p65 subunit were enhanced in pasakbumin A-treated Raw264.7 cells during Mtb infection compared to cells infected with Mtb alone (**Fig. 2A**). Our previous study demonstrated that lysophosphatidylcholine (LPC) controls intracellular Mtb survival by promoting phagosomal maturation in Mtb-infected cells, based on elevated levels of cleaved cathepsin D[29], a lysosomal aspartic protease that undergoes several proteolytic processing events to produce a mature protein at an acidic pH. To investigate whether pasakbumin A can control phagosomal maturation in Mtb-infected cells, we also detected the expression level of cleaved cathepsin D in Mtb-infected cells. As shown in **Fig. 2A**, cathepsin D was rapidly cleaved in pasakbumin-treated cells 30min after Mtb infection compared to cells infected with Mtb alone.

**Fig 2.**
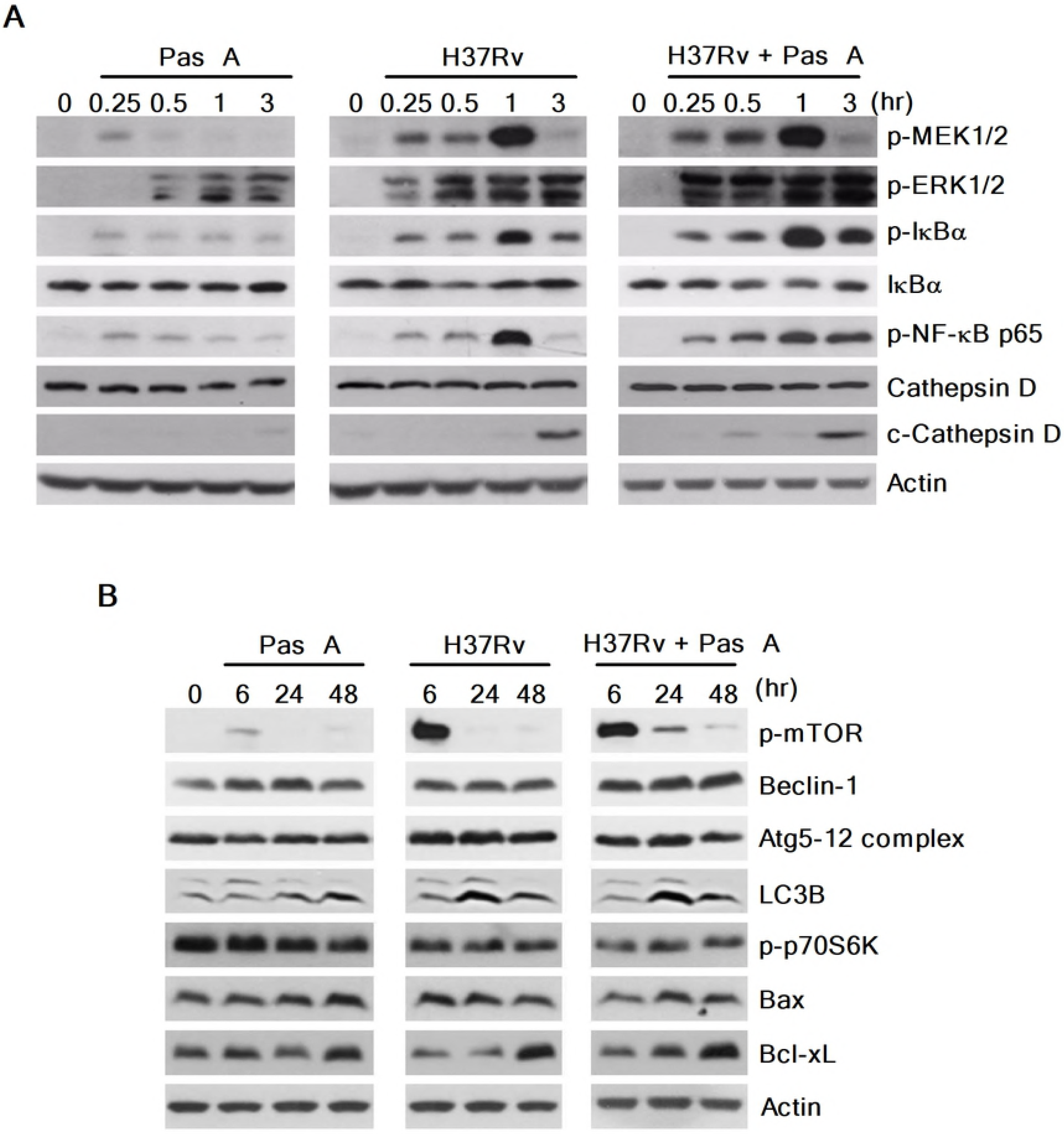
Pasakbumin A activates the NF-κB- and ERK1/2-mediated signaling pathways and induces autophagy in Mtb-infected macrophages. Raw264.7 macrophages were infected with H37Rv (at an MOI of 5) and then treated with pasakbumin A (Pas A, 10 μM) for the indicated time points. Western blot analysis showed the expression of various proteins form (A) the NF-κB and MAPK signaling pathways and (B) the expression of apoptosis- and autophagy-related proteins in pasakbumin A-treated macrophages during H37Rv infection. (A) A complete western blots are provided in **S2 Data File**. (B) Cropped membranes from different gels were used in the western blot assay.

We also demonstrated that infection with the virulent Mtb strain H37Rv induced more necrosis than the avirulent Mtb strain H37Ra in bone marrow-derived macrophages, leading to host cell death and bacterial dissemination[30]. Therefore, various cell death mechanisms are essential to host defense mechanisms by which host cells remove intracellular pathogens and pathogen-infected host cells[31]. Notably, apoptosis and autophagy play a pivotal role in the pathogenesis as well as the host defense mechanisms against Mtb infection[32]. We therefore examined whether pasakbumin A affected the induction of different cell death mechanisms, including autophagy and apoptosis. As shown **Fig. 2B**, pasakbumin A strikingly increased the phosphorylated mTOR level as well as the conversion of LC3-I to LC3-II in Mtb-infected Raw264.7 cells compared to untreated control cells. Interestingly, pasakbumin A also increased the anti-apoptotic protein Bcl-xL, whereas it decreased the pro-apoptotic protein Bax in H37Rv-infected Raw264.7 cells (**Fig. 2B**). These data demonstrated that pasakbumin A induced autophagy with activation of the ERK1/2- and NF-κB signaling pathways in Mtb-infected macrophages. Moreover, pasakbumin A blocked host cell death via apoptosis.

### Pasakbumin A regulates intracellular Mtb growth and activates the production of inflammatory mediators via the ERK1/2 signaling pathway

In Fig. 1 we show that pasakbumin A suppressed intracellular Mtb growth and enhanced the production of inflammatory mediators, including TNF-α and NO, in H37Rv-infected Raw264.7 cells. Therefore, to further demonstrate which signaling pathway inhibits intracellular Mtb growth and produces inflammatory mediators in pasakbumin A-treated macrophages during Mtb infection, we focused on the ERK1/2-mediated signaling pathway. We used the pharmacologic inhibitor U0126 to inhibit the ability of MEK1/2 to activate ERK1/2. As shown **Fig. 3A**, inhibition of ERK1/2 entirely failed to suppress intracellular Mtb growth in the presence or absence of pasakbumin A during H37Rv infection. In addition, treatment with the ERK1/2 inhibitor decreased NO and TNF-α production, whereas it slightly increased IL-10 production in pasakbumin A-treated Raw264.7 cells during H37Rv infection, similar to the effect observed in cells infected with H37Rv alone (**Fig. 3B-3D**). These results indicated that ERK1/2-mediated signaling could control intracellular Mtb growth and to produce inflammatory mediators in pasakbumin A-treated macrophages.

**Fig 3.**
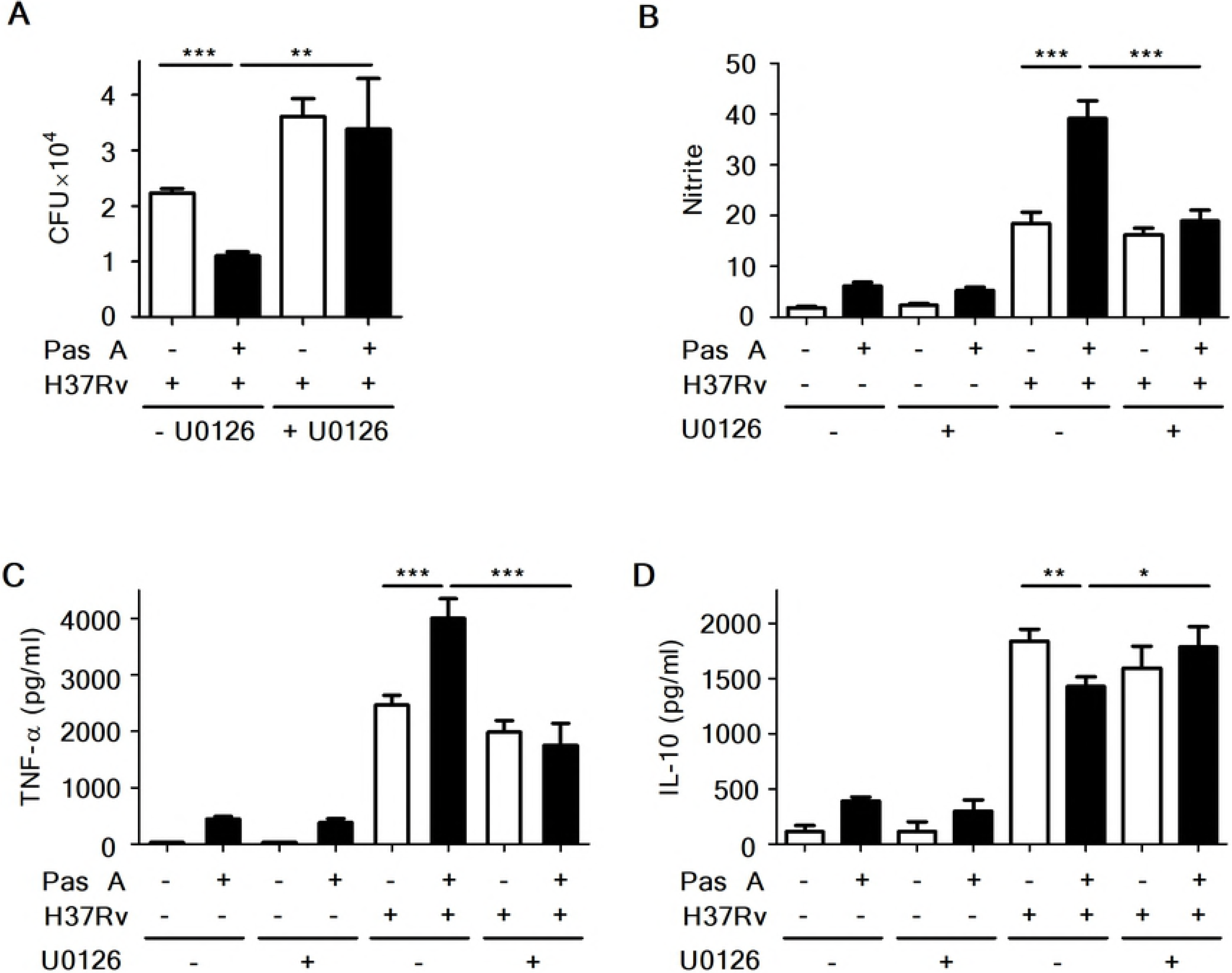
Pasakbumin A controls intracellular Mtb growth and production of inflammatory mediators through ERK-mediated signaling. Raw264.7 cells were pre-treated with U0126 (10 μM) for 1 h and then stimulated with pasakbumin A (Pas A, 10 μM) during H37Rv infection (MOI of 1 or 5). (A) Intracellular bacterial survival was determined by counting the number of CFUs 3 weeks after inoculation. (B-D) Culture supernatants were assessed for the production of (B) NO, (C) TNF-α and (D) IL-10 at 48 h. The experiments were performed in triplicate. *, *p*<0.05; **, *p*<0.01 and ***, *p*<0.001.

### ERK1/2-mediated signaling induces autophagy in pasakbumin A-treated macrophages during Mtb infection

A recent study showed that blockade of ERK1/2 inhibits *beclin-1* or *Atg5* mRNA expression in vitamin D3-treated THP-1 cells during Mtb infection, indicating that the induction of autophagy depends on ERK1/2-mediated signaling during Mtb infection[33]. To assess whether pasakbumin A-induced autophagy is triggered by ERK1/2-mediated signaling in Mtb-infected macrophages, we examined the induction of autophagy in pasakbumin A-treated Raw264.7 cells in the presence or absence of U0126. Treatment with U0126 diminished ERK1/2 and IκBα phosphorylation in pasakbumin A-treated Raw264.7 cells during H37Rv infection, similar to those found in Mtb-infected Raw264.7 cells without U0126 (**Fig. 4A**). Treatment with pasakbumin A enhanced the number of endogenous LC3-positive puncta, which indicated autophagosomes, in Mtb-infected Raw264.7 cells compared with those infected with H37Rv without pasakbumin A (**Fig. 4B**). However, the number of endogenous LC3-positive puncta was significantly reduced by the ERK1/2 inhibitor U0126 in pasakbumin A-treated Raw264.7 cells during H37Rv infection, similar to those infected with H37Rv alone (**Fig. 4B**). These results indicated that ERK1/2-mediated signaling was important not only for autophagy induction but also for the production of inflammatory mediators after pasakbumin A treatment.

**Fig 4.**
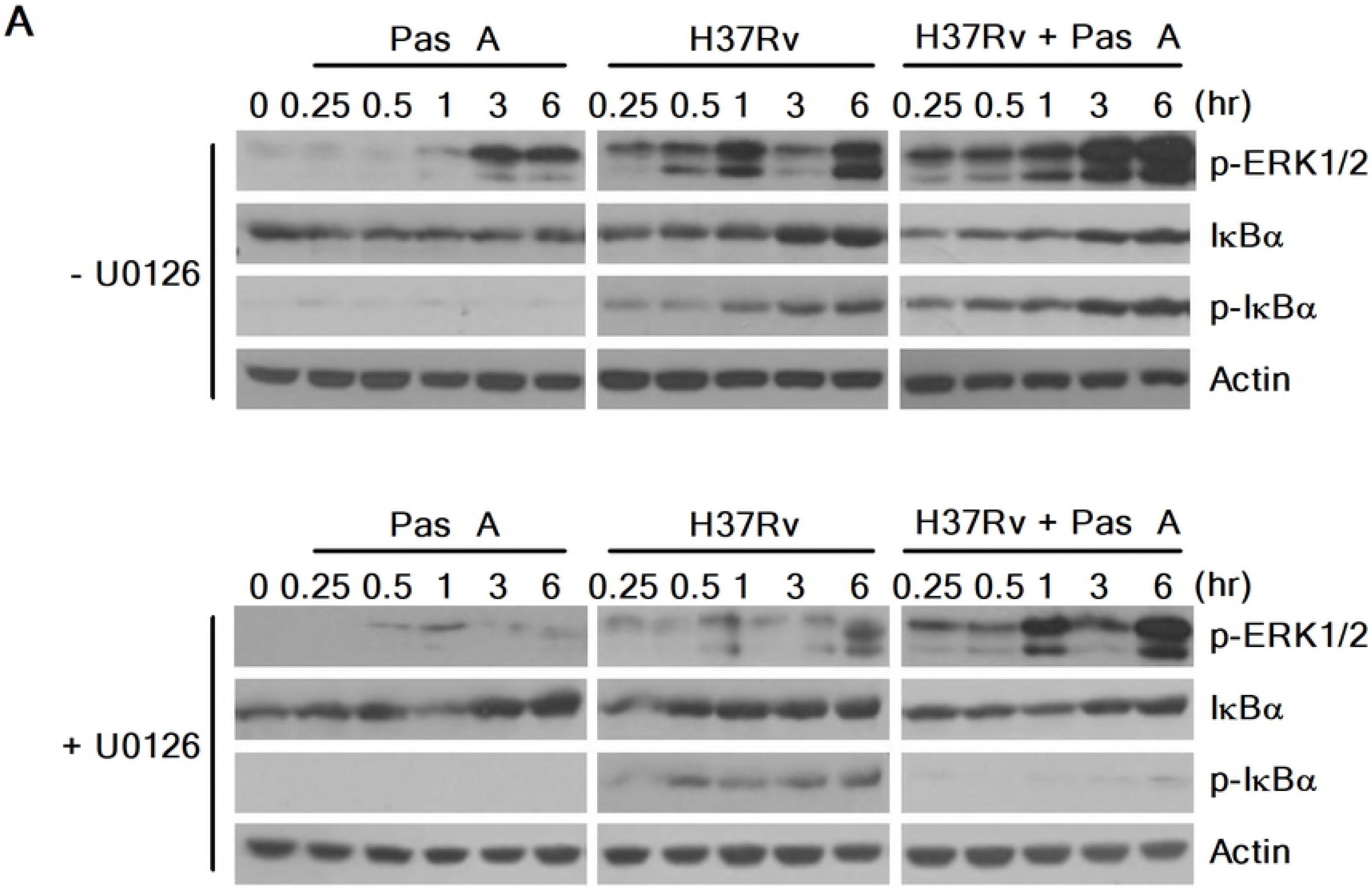
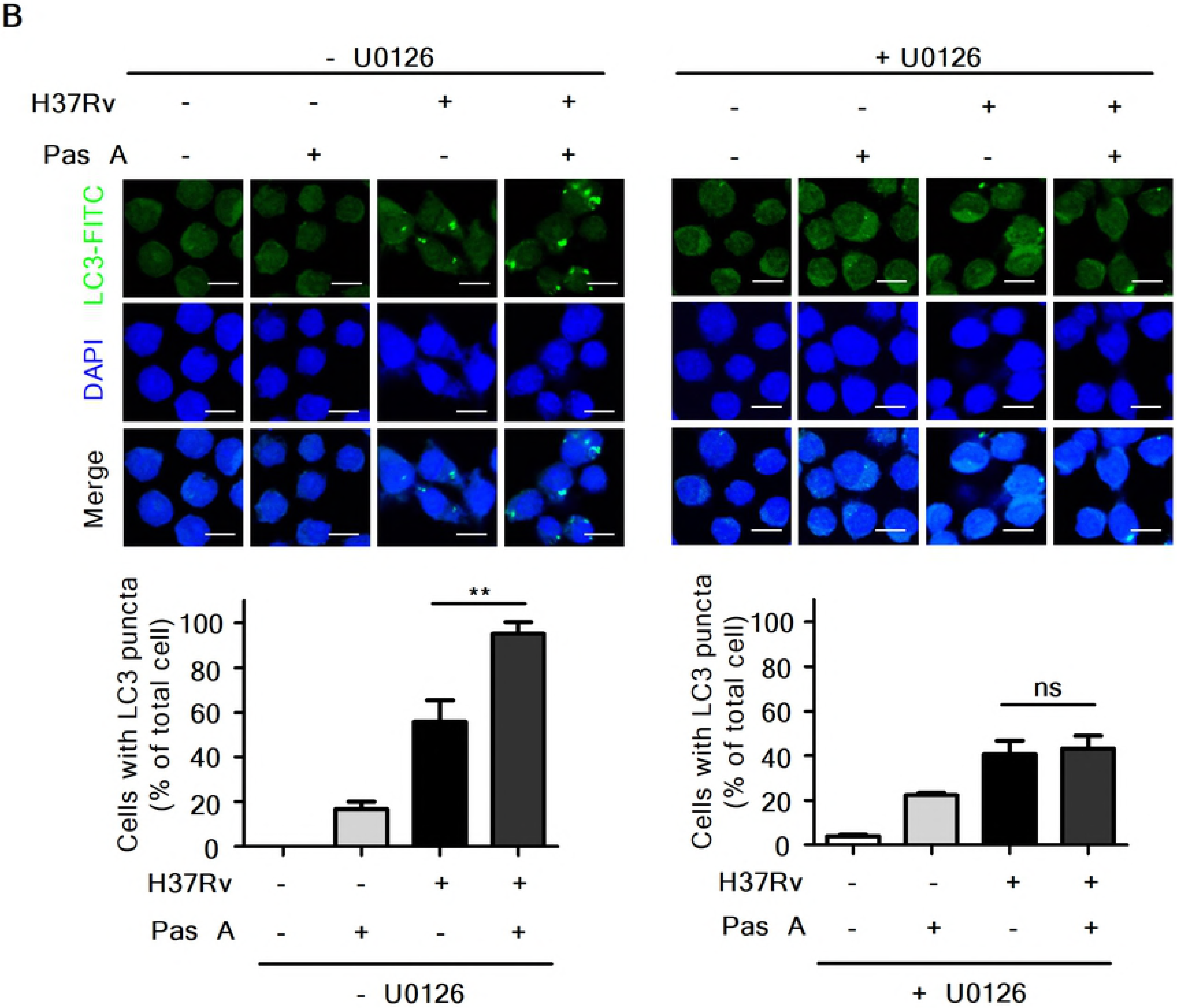
Pasakbumin A induces autophagy by activating ERK1/2-mediated signaling. Raw264.7 cells were pre-treated with U0126 (10 μM) for 1 h and then stimulated with pasakbumin A (Pas A, 10 μM) during H37Rv infection (MOI of 1 or 5). (A) The changes in the phosphorylated and total protein levels of ERK1/2 and iKBa were assessed using western blot analysis. Cropped membranes from different gels were used in the western blot assay. (B) Immunofluorescence staining of LC3 in Mtb-infected cells treated as described above. The number of LC3-positive puncta was counted under a microscope, and the percentage of cells containing LC3-positive puncta relative to the total cell number was calculated. Statistical significance is indicated as *, *p*<0.05 and ns, not significant (*p*>0.05).

### Combined treatment with an anti-TB drug improves mycobactericidal activity of pasakbumin A in Mtb-infected macrophages

Recently, a published report showed that treatment with the anti-TB drugs INH and RMP increased cellular and mitochondrial reactive oxygen species (ROS), leading to autophagy activation in Mtb-infected macrophages. These findings indicated that antibiotic-induced autophagy plays an important role in chemotherapy against Mtb infection[34]. We next questioned whether combined treatment with an anti-TB drug is more efficacious than treatment with pasakbumin A alone for infection with Mtb. To investigate this, we assessed intracellular bacterial growth in Mtb-infected Raw264.7 cells in the presence or absence of the anti-TB drug RMP. As shown **Fig. 5A**, treatment with RMP suppressed intracellular bacterial growth in a dose-dependent manner. Notably, RMP at a dose with significant antimicrobial activities (0.5 μg/ml for RMP) significantly inhibited intracellular bacterial survival in pasakbumin A-treated cells during Mtb infection compared to that of Mtb-infected cells (**Fig. 5A**). In addition, TNF-α secretion was irregularly increased by RMP in a dose-dependent manner; however, more was produced in Mtb-infected cells treated with pasakbumin A and RMP than Mtb-infected cells treated with RMP alone (**Fig. 5B**). In contrast, combined treatment of pasakbumin A and RMP showed stronger decreases in IL-10 secretion than that of cells treated with pasakbumin A in the absence of RMP during Mtb infection (**Fig. 5C**). There was no significant difference in NO production after RMP treatment in pasakbumin A-treated cells (**Fig. 5D**). These data suggested that the combination of pasakbumin A with an anti-TB drug effectively suppressed intracellular Mtb growth by promoting the production of pro-inflammatory cytokine and blocking the production of anti-inflammatory cytokine in macrophages.

**Fig 5.**
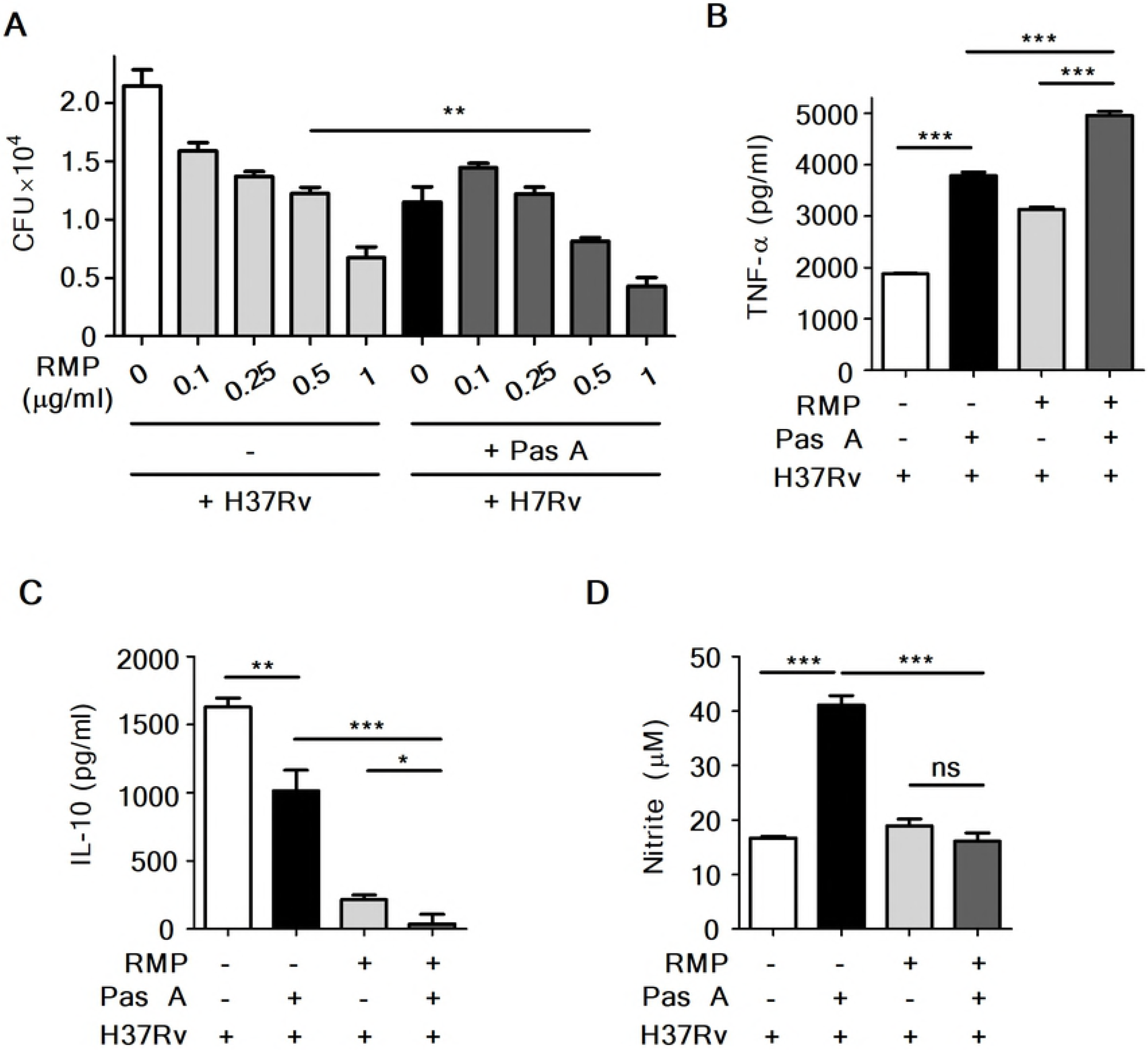
Combination treatment with pasakbumin A and an anti-TB drug improves the antibacterial immunity of Mtb-infected macrophages. Raw264.7 macrophages were infected with H37Rv (at MOI of 1 or 5) for 4 h and treated with pasakbumin A (Pas A, 10 μM) alone or Pas A combined with an anti-TB drug, rifampicin (RMP), for 48 h in a dose-dependent manner. (A) Intracellular bacterial survival was determined by counting the number of CFUs at 3-weeks after inoculation. (B, C) Culture supernatants were assessed for the production of (B) TNF-α and (C) IL-10 with ELISA at 48 h. (D) NO production was detected in cell culture supernatants. Statistical significance is indicated as *, *p*<0.05; **, *p*<0.01, ***, *p*<0.001 and ns, not significant (*p*<0.05).

### RMP strongly accelerates pasakbumin A-induced autophagy through ERK1/2-mediated signaling in Mtb-infected macrophages

Our present data showed that pasakbumin A induced autophagy via ERK1/2-mediated signaling, and combined treatment with RMP substantially reduced intracellular bacterial growth in Mtb-infected macrophages. Thus, we investigated the cellular mechanism by which combined treatment of pasakbumin A and RMP enhances autophagy to control intracellular Mtb growth in macrophages. Phosphorylated levels of ERK1/2 and IκBα rapidly increased after the combined treatment with pasakbumin A and RMP in Mtb-infected cells compared with those treated with pasakbumin A alone (**Fig. 6A**). We also found that combined treatment of pasakbumin A and RMP strongly induced the conversion of LC3-I to LC3-II in Mtb-infected cells (**Fig. 6A**). To further confirm this observation, we detected endogenous LC3 by immunofluorescence staining. After combined treatment of pasakbumin A and RMP, the percentage of FITC-LC3-positive cells with punctate staining was greater than that of Mtb-infected cells treated with pasakbumin A alone (**Fig. 6B**). These data suggest that pasakbumin A may be an effective adjuvant for enhancing the host-directed immunotherapy against tuberculosis.

**Fig 6.**
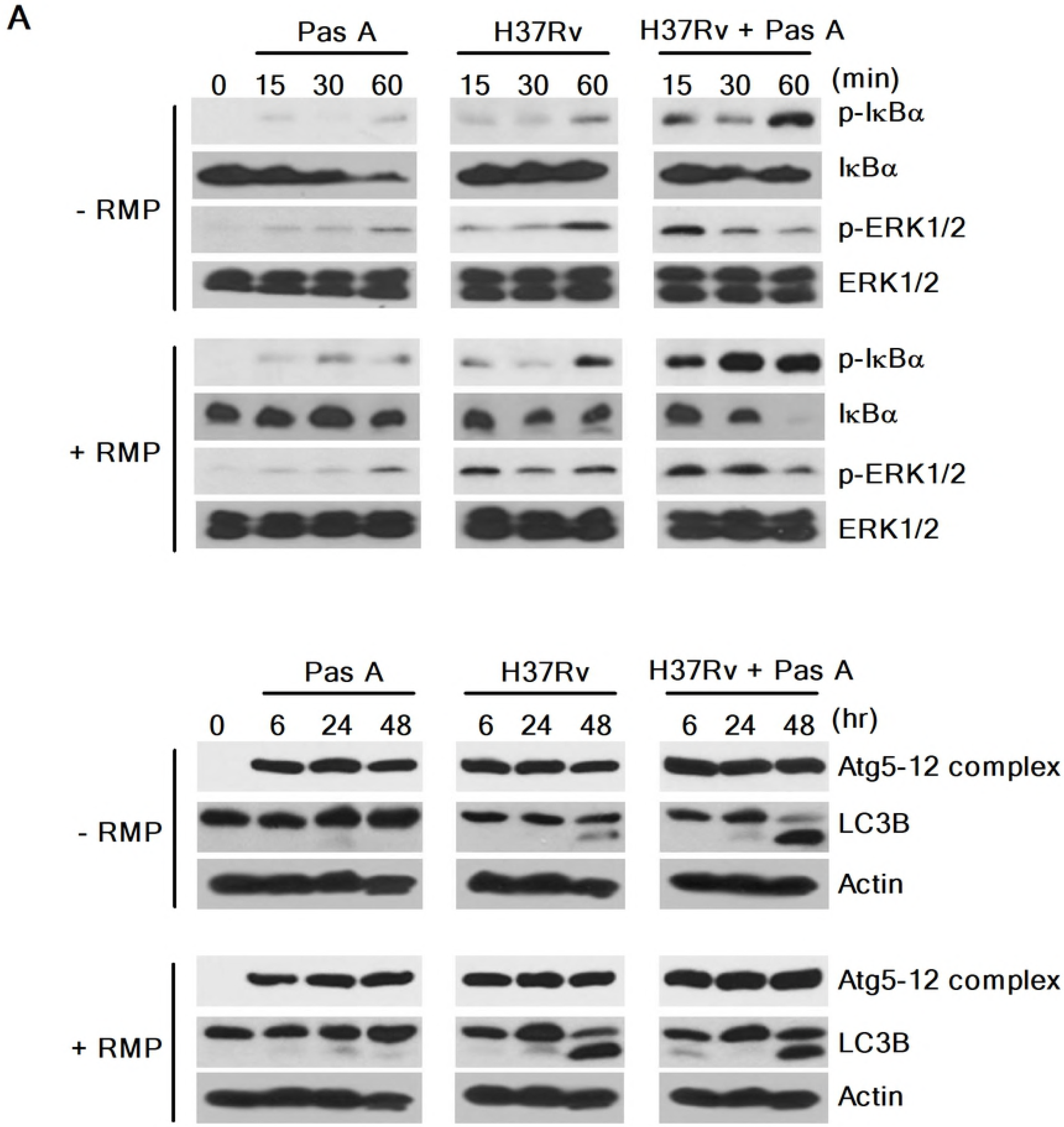
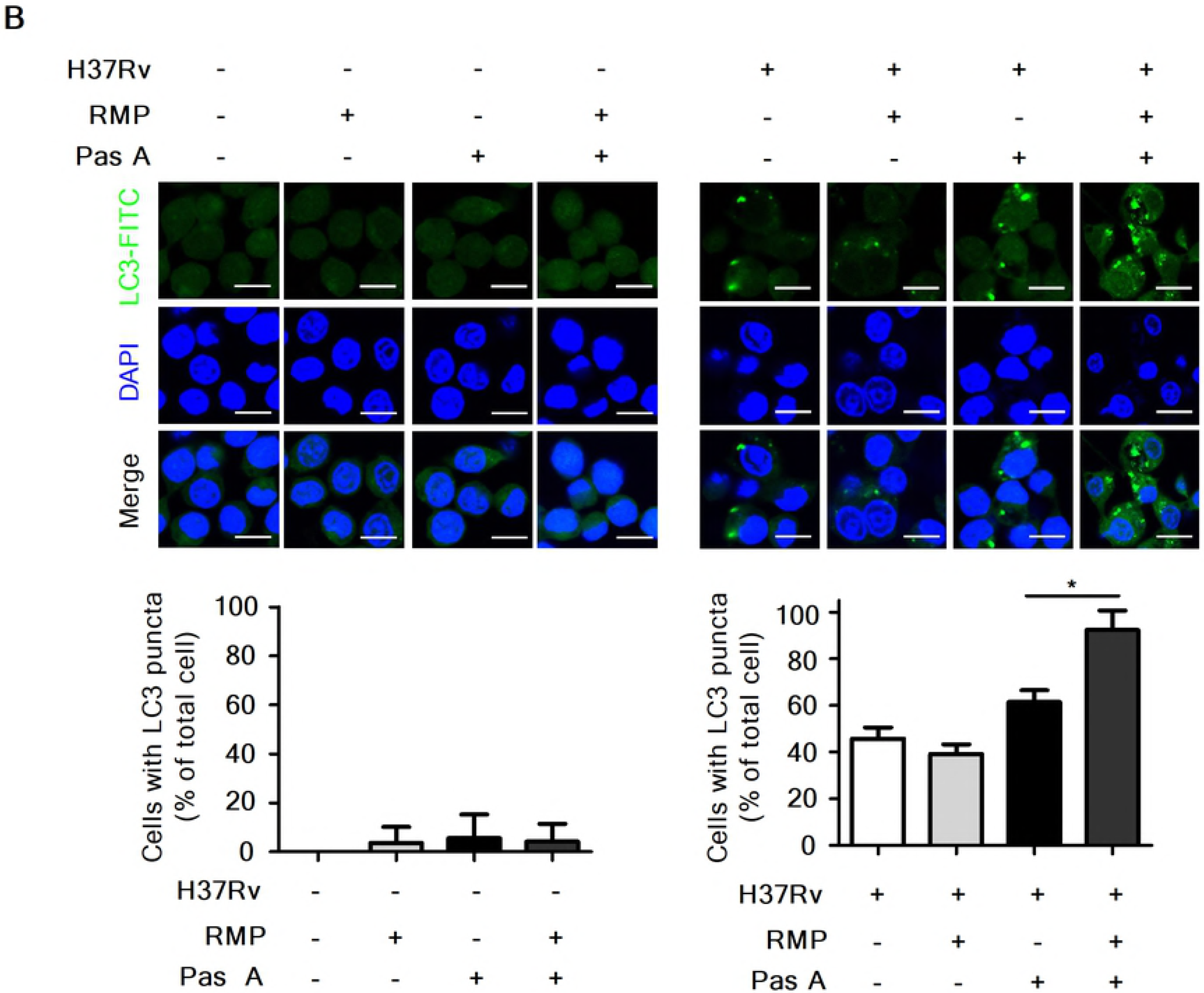
Treatment of pasakbumin A combined with RMP accelerates autophagy and the activation of NF-κB- and ERK1/2-mediated signaling compared to that of cells treated with pasakbumin A alone during Mtb infection. (A) Raw264.7 macrophages were infected with H37Rv (at a MOI of 5), and then treated with pasakbumin A (Pas A, 10 μM) for the indicated time points. (A) The expression levels of autophagy-related proteins as well as phosphorylated and total protein for various components of the NF-κB and MAPK signaling pathways were examined by the western blot assay. Cropped membranes from different gels were used in western blot assay. (B) The LC3-positive macrophages after treatment with Pas A with or without RMP during Mtb infection were detected via immunofluorescence staining. The number of LC3-positive puncta was counted under a microscope, and the percentage of cells containing LC3-positive puncta relative to the total cell number was calculated. Statistical significance is indicated as *, *p*<0.05.D

## Discussion

TB, which is caused by Mtb, remains a major infectious disease despite the advances made in treatment and management[1]. TB can be effectively treated with a multidrug regimen of first-line drugs, including INH, RMP, PZA, EMB and streptomycin (SM), for at least 6 months[35]. However, primary therapy often fails to cure TB for several reasons, including patient non-compliance, inappropriate drug levels, drug shortages and a number of other factors[36]. In this context, Menzies *et al* showed that TB patients non-compliances was the main reason for treatment failure because patients did not take prescribed medication according to the records of Montreal Chest Hospital from 1987-1988[37].

The multidrug regimen of anti-TB drugs has also been associated with an increased incidence of adverse effects that cause mild to severe host damage; these effects may cause discontinuation of the treatment due to the poor health and immunity of TB patients[20]. Common adverse effects are dizziness, muscular twitching, loss of vision or hearing, hemolytic anemia, acute renal failure, hepatotoxicity and thrombocytopenia[38, 39]. Recently, Yakar *et al* showed that thrombocytopenia is induced by INH and RMP, and patients had no further thrombocytopenia development without INH and RMP during the therapeutic period[40]. Furthermore, many other studies have demonstrated the regulatory effects of RMP on the immune response, including phagocytosis, antibody production, T cell differentiation and delayed hypersensitivity[41]. Due to these problems, there is an urgent need to identify novel anti-TB agents.

*E. longifolia* (known as tongkat ali) is a commonly distributed and popular traditional herbal medicine in Southeast Asia and Indo-China used to treat various illnesses, including sexual dysfunction, fever, malaria, ulcers, high blood pressure, TB, and diarrhea[26]. The plant portions of *E. longifolia* have a variety of bioactive compounds, such as quassinoids, alkaloids and squalene derivatives[42]. Tada *et al.* isolated four quassinoids, pasakbumin-A, -B, -C, and -D, from the roots and showed that pasakbumin-A and -B have anti-ulcer activity[26]. Many studies have also shown that pasakbumin-A, -B, -C, and -D have strong cytotoxicity toward human lung cancer (A549) and human breast cancer (MCF-7) cell lines[43, 44]. Farouk and Benafri demonstrated that the aqueous extracts of leaves exhibit antibacterial activity against *Staphylococcus aureus* and *Serratia marscesens*[27], however, antibacterial activity against other pathogenic microorganisms has not been determined. In the present study, pasakbumin A extracted from *E. longifolia* was shown to have anti-TB activity against the virulent Mtb strain H37Rv in a Raw264.7 macrophage cell line. As a result, our hypothesis of the anti-TB activity of pasakbumin A appears to be accurate because pasakbumin A inhibited intracellular Mtb growth in mouse macrophages. However, pasakbumin A showed little cytotoxic activity in the present study on Mtb and mouse macrophages. In contrast, in H37Rv-infected macrophages, pasakbumin A protected the host cells from Mtb-induced apoptotic cell death. Our results suggest that pasakbumin A can be developed as a safe and continuously available therapeutic drug to protect hosts from cell death and interfere with the growth of intracellular Mtb.

Autophagy is a lysosomal self-degradation process for cellular homeostasis and functions as an innate defense mechanism during Mtb infection. Gutierrez *et al.* first showed that induction of autophagy by starvation or treatment with the mTOR inhibitor rapamycin increases co-localization of Mtb with LC3 and beclin-1 and delivers Mtb to phagolysosomes, suggesting that autophagy plays an important role as a defense mechanism against TB[45]. Our results also suggested that activation of autophagy induced by pasakbumin A may play a central role in its antibacterial effects.

Many studies have demonstrated that several antibiotics can induce or suppress autophagy. Kim *et al.* showed that INH and PZA promoted autophagy activation and phagosomal maturation in Mtb-infected host cells, suggesting that host autophagy plays an important role in host protective responses during antibiotic chemotherapy against TB[34]. In this context, we observed that the combined treatment of pasakbumin A and the anti-TB drug RMP resulted in increased intracellular LC3 distribution and phosphorylated ERK1/2 and IκBα in Mtb-infected cells compared to cells infected Mtb alone. These results reveal that the combined treatment of pasakbumin A with an anti-TB drug resulted in effective antimicrobial activity against Mtb infection by inducing autophagy and that pasakbumin A also protects host cells from apoptotic cell death during Mtb infection.

In conclusion, we report that pasakbumin A inhibited intracellular Mtb growth in mouse macrophages by activating autophagy through the ERK1/2-mediated signaling pathway. This compound also increased NO and pro-inflammatory cytokine levels via the ERK1/2- and NF-κB-mediated signaling pathways in Mtb-infected cells but not the anti-inflammatory cytokine IL-10. Treatment with the anti-TB drug RMP in combination with pasakbumin A significantly suppressed intracellular Mtb growth by promoting autophagy and enhancing TNF-α production via the ERK1/2-mediated signaling pathway. Overall, understanding the molecular mechanisms by which pasakbumin A fights Mtb infection will provide insights into the development of novel therapeutic anti-TB drugs and potential HDT strategies that modulate the host immune response against Mtb infection.

## Materials and Methods

### Cell culture

The mouse macrophage cell line, Raw264.7, was purchased from ATCC (ATCC, Rockville, MD, USA). Cells were maintained in RPMI-1640 (Cellgro, Herndon, VA, USA) supplemented with 10% fetal bovine serum (FBS, Atlas Biologicals, Fort Collins, CO) and 1% penicillin/streptomycin (Corning Incorporated, Corning, NY, USA). Cells were cultured in a standard cell culture incubator at 37°C with an atmosphere of 5% CO_2_ and 95% air.

### Reagents

Pasakbumin A was isolated from *E. longifolia* roots as previously described[44]. RMP and INH (isonicotinic acid hydrazide) were purchased from Tokyo Chemical Industry Co. Ltd. (TCI Co. Ltd., Tokyo, Japan). Other reagents were as follows: U0126 (Cell Signaling, Danvers, MA, USA), 3-methyladenine (3-MA, Sigma-Aldrich, St. Louis, MO, USA), and Bay11-7082 (Cayman Chemical, MI, USA).

### Bacterial strains and culture conditions

The *M. tuberculosis* strains H37Rv and H37Ra were used in this study. Mtb was grown in Middlebrook 7H9 broth (Difco Laboratories, USA) supplemented with 10% ADC (5% bovine albumin, 2% dextrose, 0.03% catalase, 0.85% sodium chloride) and 0.2% glycerol at 37°C. After 3 weeks of culture, Mtb was harvested, adjusted to 1 × 10^7^ bacteria/200 κl stock solution, aliquoted, and maintained at −70°C until used.

### Enzyme-linked immunosorbent assay (ELISA)

Cultured supernatants were collected and tested for TNF-α and IL-10 production using an ELISA kit (Peprotech, NJ, USA) according to the manufacturer’s instructions. Samples were read at 450 nm using a microplate reader (Biotek Instruments Inc., Winooski, VT, USA).

### Nitric oxide (NO) detection assay

NO production was measured using a nitric oxide detection kit (Intron Bio-technology Inc., Kyungki-Do, Korea) according to the manufacturer’s procedure. NO detection was performed as previously described[29]. Briefly, culture supernatants were collected and mixed with N1 buffer (sulfanilamide in the reaction buffer) for 10 min at room temperature. After 10 min, the mixture was combined with N2 buffer (naphthylethylenediamine in the stabilizer buffer) for 10 min at room temperature, and absorbance was measured between 520-560 nm using a plate reader. NO production was calculated from a standard curve with nitrite standard solution.

### Western blot assay

For cell lysates, the cells were lysed with lysis buffer containing complete protease inhibitor cocktail (Calbiochem, San Diego, CA, USA). Western blotting was performed as previously described[46]. The acrylamide percentage of the SDS-PAGE gel was determined by the size of the target protein in the sample. The membranes were stripped or cropped to detect multiple proteins. Anti-NOS2, anti-IκBα, anti-NF-κB p65, anti-cathepsin D, anti-Bax, anti-Bcl-xL and anti-β-actin were purchased from Santa Cruz Biotechnology (Santa Cruz, CA, USA). Anti-phospho-MEK1/2, anti-phospho-ERK1/2, anti-phospho-IκBα, anti-phospho-mTOR, anti-beclin-1, anti-Atg5-12 complex, anti-LC3B and anti-phospho-p70S6K were purchased from Cell Signaling (Danvers, MA, USA).

### Colony-forming unit (CFU) assay

Mtb-infected macrophages were lysed with 0.1% saponin (Sigma-Aldrich, Dorset, UK) for 10 min at 37°C with 5% CO_2_ and then serially diluted in Middlebrook 7H9 broth. Next, samples were plated on Middlebrook 7H10 agar in triplicate and incubated at 37°C for 21 days. CFUs were counted 21 days after incubation.

### Immunofluorescence analysis

Immunofluorescence analysis was performed as previously described[47]. Briefly, cells were seeded onto cover-slides and treated as described above. Cells were washed in PBS, fixed in 4% paraformaldehyde overnight, and permeabilized in 0.1% Triton X-100 for 15 min. Slides were incubated with the primary antibodies (anti-LC3B, MBL, Nagoya, Japan) at room temperature for 2 h and washed 3 times with PBS for 5 min. Then, the slides were incubated with FITC anti-rabbit secondary antibody (Jackson Immunoresearch, West Grove, USA) at room temperature for 1.5 h, and nuclei were counterstained with 4’-6-diamidino-2-phenylindole (DAPI, Sigma-Aldrich, MO, USA) for 5 min. Cover-slides were mounted in Fluoromount-G™ and examined by confocal microscopy (FV1000 SPD, Olympus, Tokyo, Japan).

### Cell viability assay

Raw264.7 cells were infected with H37Rv at a MOI of 1 for 4 h and then treated with pasakbumin A for 72 h. Cells were suspended and stained with trypan blue solution (Sigma-Aldrich, MO, USA), which selectively stains dead cells. The cell number was determined by a hemocytometer.

### Statistical analysis

The results are presented as the mean±SD of triplicate experiments. Statistical significance was analyzed using Student’s t-test and one-way ANOVA followed by Tukey’s post hoc test for multiple comparisons. The data were graphed and analyzed using GraphPad Prism software (GraphPad Software, La Jolla, CA, USA). Values of *p*<0.05 were considered significant. Statistical significance is indicated as *, *p*<0.05; **, *p*<0.01; ***, *p*<0.001; and ns, not significant (*p*>0.05).

## Acknowledgments

This research was supported by a grant from the Korea Health Technology R&D Project through the Korea Health Industry Development Institute (KHIDI), which is funded by the Ministry of Health & Welfare of the Republic of Korea (HI15C0450), and the Basic Science Research Program through the National Research Foundation of Korea (NRF), which is funded by the Ministry of Education, Science and Technology (2017R1A6A3A11032251, 2018R1D1A1B07049097). Funding was also provided by a Research Grant from Kangwon National University in 2016.

## Supporting information

**S1 Data File. Complete Western blots**.

**S2 Data File. Complete Western blots**.

